# Copy number heterogeneity, large origin tandem repeats, and interspecies recombination in HHV-6A and HHV-6B reference strains

**DOI:** 10.1101/193805

**Authors:** Alexander L. Greninger, Pavitra Roychoudhury, Negar Makhsous, Derek Hanson, Jill Chase, Gerhard Krueger, Hong Xie, Meei-Li Huang, Lindsay Saunders, Dharam Ablashi, David M. Koelle, Linda Cook, Keith R. Jerome

## Abstract

Quantitative PCR is the diagnostic pillar for clinical virology testing, and reference materials are necessary for accurate, comparable quantitation between clinical laboratories. Accurate quantitation of HHV-6 is important for detection of viral reactivation and inherited chromosomally integrated HHV-6 in immunocompromised patients. Reference materials in clinical virology commonly consist of laboratory-adapted viral strains that may be affected by the culture process. We performed next-generation sequencing to make relative copy number measurements at single nucleotide resolution of eight candidate HHV-6A and seven HHV-6B reference strains and DNA materials from the HHV-6 Foundation and Advanced Biotechnologies. 11 of 17 (65%) HHV6 candidate reference materials showed multiple copies of the origin of replication upstream of the U41 gene by next-generation sequencing. These large tandem repeats arose independently in culture-adapted HHV-6A and HHV-6B strains, measuring 1254 bp and 983 bp, respectively. Copy number measured between 4-10X copies relative to the rest of the genome. We also report the first interspecies recombinant HHV-6 strain with a HHV-6A GS backbone and >5.5kb region from HHV-6B Z29 from U41-U43 that covered the origin tandem repeat. Specific HHV-6A reference strains demonstrated duplication of regions at UL1/UL2, U87, and U89, as well as deletion in the U12-U24 region and U94/95 genes. HHV-6 strains derived from cord blood mononuclear cells from different labs on different continents revealed no copy number differences throughout the viral genome. These data indicate large origin tandem duplications are an adaptation of both HHV-6A and HHV-6B in culture and show interspecies recombination is possible within the *Betaherpesvirinae.*

**Importance:** Anything in science that needs to be quantitated requires a standard unit of measurement. This includes viruses, for which quantitation increasingly determines definitions of pathology and guidelines for treatment. However, the act of making standard or reference material in virology can alter its very usefulness through genomic duplications, insertions, and rearrangements. We used deep sequencing to examine candidate reference strains for HHV-6, a ubiquitous human virus that can reactivate in the immunocompromised population and is integrated into the human genome in every cell of the body for 1% of people worldwide. We found large tandem repeats in the origin of replication for both HHV-6A and HHV-6B that are selected for in culture. We also found the first interspecies recombinant between HHV-6A and HHV-6B, a phenomenon that is well-known in alphaherpesviruses but to date has not been seen in betaherpesviruses. These data critically inform HHV-6 biology and the standard selection process.

## Introduction

Human herpesvirus 6A and 6B (HHV-6) are ubiquitous human viruses with human exposure levels >90% by the age of 2 years old as measured by serological assays performed worldwide (1, 2). Both HHV-6A and HHV-6B establish chronic infections in the majority of infected individuals, leading to asymptomatic persistent viral shedding (3, 4). Exanthema subitum is the most common HHV-6 related infection seen after a primary exposure in 6 month to 3-year old children. Less frequently the virus can result in seizures, gastrointestinal and respiratory symptoms, thrombocytopenia, hepatitis, colitis, and CNS infections. Additionally, both HHV-6A and HHV-6B have been shown to integrate into host chromosomes in the telomere regions and be passed from parents to their children as inherited chromosomally integrated HHV-6 (iciHHV-6). The potential to measure HHV-6 from genomic DNA from these patients, as well as possible reactivation from the integrated HHV-6 make the diagnosis of HHV-6 infection from serum or plasma viral load testing challenging.

Accurate and sensitive real-time PCR assays that detect and quantify HHV-6 are critical to diagnosis and monitoring of the variety of manifestations of HHV-6 associated disease. Recently several quantitative cut-offs have been proposed that are associated with end-organ disease or iciHHV-6 status (5–7). There have been only a few limited studies comparing PCR methods between clinical labs (8–10). In addition, newer studies often utilize commercial reference laboratories or commercial reagents that in general do not make their primer/probe locations known. Of published studies where the PCR locations are described, many areas of the genome have been used, with no consistent location chosen. A review of the PCR methods used in 46 recent published papers (years 2014-2017) revealed the use of 17 different primer sets at multiple locations throughout the genome (at U6, 12, 13, 22, 27, 31, 32, 38, 41, 57, 65, 66, 67, 69, 90, 95, 100), with only the U31 and U65-66 primers used more than twice. Not surprisingly, this lack of consistency has contributed to a significant lack of consistency in test results as measured in cross-lab proficiency testing where quantitative differences as high as 4 logs have been seen (10, 11). These studies hint that between-lab results may be improved if diagnostic testing was done with a more limited number of high-performing primer sets. Finally, effective primer designs have been significantly limited by the lack of available DNA sequences.

Previous studies have identified the critical role that standardized materials play in the ability to establish clinical viral load cut-offs, establish assay sensitivity, and compare results between laboratories (10, 12–16). Efforts are currently underway at the National Institute for Biological Standards and Controls (NIBSC) in the United Kingdom to prepare WHO international standard material for both HHV-6A and 6B. However, since this effort utilizes cultured HHV-6 reference strains, it is unknown how well the materials utilized will reflect sequences in clinical isolates. Given the wide range of primer set locations used in labs, it is critical that the entire genome of any reference materials be studied

HHV-6A and -6B genomes have approximately 90% nucleotide identity to each other and about 50% similarity with the closest-related human betaherpesvirus, HHV-7. The genome is approximately 160-170 kb and contains many of the gene and regulatory elements present in the genome of other betaherpesviruses. Several prototypic strains of HHV-6 have been identified and utilized including GS, U1102, SIE, LHV, Z29, and HST. Until very recently, there were fewer than 200 HHV-6 sequences in Genbank, including only 3 complete genomes. Recently, two large scale efforts have sequenced more than 150 near-full length HHV-6B genomes from four continents (17, 18).

In an effort to determine whether the available cultured “reference” strains have undergone significant changes during culture similar to that seen with the recently produced WHO BK and JC strains, we obtained 15 strains from the HHV-6 Foundation repository and used shotgun sequencing to obtain full-length genomes and estimate of copy number (19, 20). We found 9 of the 15 strains had high-copy tandem repeat amplifications in the origin of replication, including the first described origin amplification in an HHV-6A strain. We also describe the first HHV-6A/HHV-6B interspecies recombinant. Other HHV-6 reference materials had multiple loci with copy number variation of up to 20X. These duplications, deletions, and rearrangements may impact the utility of these strains for the production of standard materials for PCR testing. Changes in the genome of these strains in culture may have impact on the results of current and future studies utilizing these materials.

## Materials and Methods

### HHV-6 reference strains and DNA materials

HHV-6 culture isolates were obtained from the HHV-6 Foundation. The original HHV-6 isolate GS belongs to HHV-6A and was first isolated at the National Cancer Institute, NIH in 1986 from an AIDS patient. The HHV-6A GS strain was grown in HSB2, which is a human T-cell leukemic line derived from the peripheral blood of a child. The GS early passage isolate obtained from the HHV-6 Foundation is a low-passaged HHV-6A GS isolate that was only passaged 4 times in cord blood mononuclear cells (CBMC). The GS early passage isolate was sequenced in 2013 after a brief expansion in CBMC and sequencing reads were obtained from L. Flamand (21). The HHV-6A DA strain was isolated at the NCI from a patient with chronic fatigue syndrome and was grown in the HSB2 cell line. The HHV-6A CO strains CO1, CO2, CO3, CO4, CO7 were isolated from patients with collagen vascular diseases including systemic lupus erythematosus, atypical polyclonal lymphoproliferation, rheumatoid arthritis, and unclassified collagen vascular disease (22). The HHV-6A SIE strain was isolated from an HIV-positive leukemia patient from the Ivory Coast and grown in PHA-stimulated CBMCs. The HHV-6B strains HST, KYO, ENO, and NAK were isolated from Japanese patients with exanthema subitum in 1988 (23). The HHV-6B MAR strain was originally obtained from an HIV-negative child born to an HIV-positive mother, and has been cultured in CBMCs (24). HHV-6B Z29 strain was originally isolated from an AIDS patients from Zaire and obtained from the HHV-6 Foundation stock deposited at the NIH AIDS repository and grown in the SupT1 cell line. Secondary HHV-6 standards comprising quantitated viral DNA from HHV-6A GS strain (08-945-250) and HHV-6B Z29 strain (08-923-00) were purchased from Advanced Biotechnologies Incorporated. Strains sequenced in this study are available in Table 1.

**Table 1.**
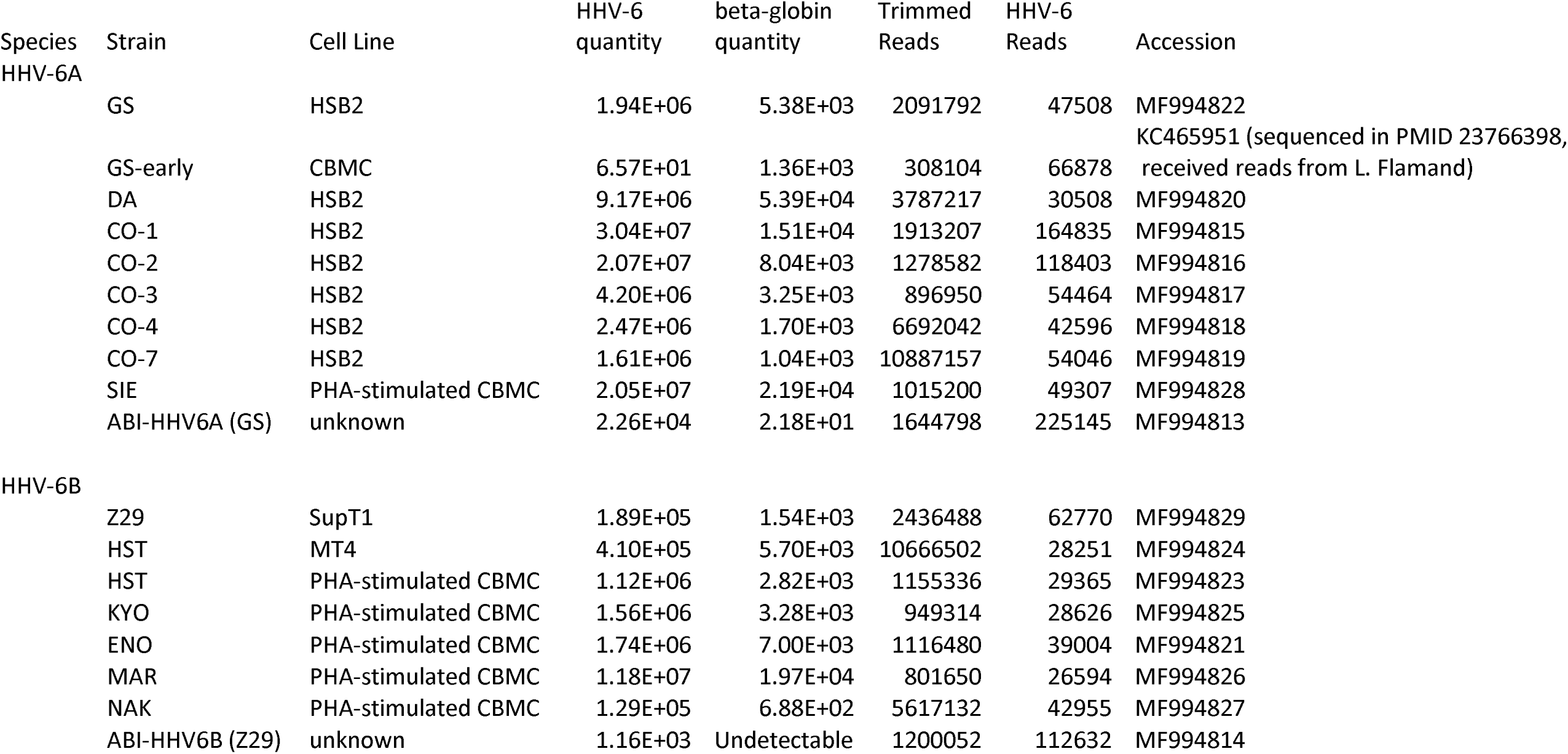
Strains sequenced with qPCR results and Sequencing

### Illumina Sequencing Library Preparation

DNA was extracted from culture isolates using the Zymo Viral DNA kit. DNA-sequencing libraries were prepared from 50 ng of genomic DNA using quarter-volumes of the Kapa HyperPrep kit with 7 minutes of fragmentation time and 12 cycles of dual-indexed Truseq-adapter PCR (18). Libraries were sequenced on 2x300bp, 1x190bp, and/or 1x192bp runs on an Illumina MiSeq. Sequences were quality- and adapter-trimmed, de novo assembled, and contigs were aligned to reference HHV-6A (NC_001664) and HHV-6B (NC_000898) genomes and visualized using Geneious v9.1. Read mapping for copy number analysis was performed using the Geneious read mapper with 10% allowed gaps per read, word length of 18, 20% maximum mismatches per read, and with structural variant, insertion, and gap finding allowed.

### qPCR confirmation

Quantitative PCR to estimate HHV-6 and beta-globin copy number in Table 1 was performed using UL67-directed directed 5R primers (GTT AGG ATA TAC CGA TGT GCG TGA T/ FAM-TCC GAA ACA ACT GTC TGA CTG GCA AAA-TAMRA/TAC AGA TAC GGA GGC AAT AGA TTT G) (25) and beta-globin primers (TGA AGG CTC ATG GCA AGA AA/ FAM-TCC AGG TGA GCC AGG CCA TCA CTA-TAMRA/ GCT CAC TCA GTG TGG CAA AGG) respectively. Briefly each 30 ul PCR reaction contained 15 ul of 2x QuantiTect Multiplex PCR NoROX Master Mix (Qiagen), 830 nM each primer, 250 nM probe, 0.267 ul Rox (Invitrogen), and 0.03 units UNG (Epicentre). EXO internal control, including template, primers and probe, was spiked into each PCR reaction to monitor PCR inhibition. QuantStudio 7 Flex Real-Time PCR system was used to perform PCR and signal detection. The PCR thermocycling conditions are as following: 50°C for 2 minutes, 95°C for 15 minutes and followed by 45 cycles of 94°C for 1 minute and 60°C for 1 minute.

Quantitive PCR to estimate relative copy number between the origin of replication and HHV-6 U32 locus was performed in 20uL reactions using the SsoAdvanced Universal SYBR Green SuperMix. Ten-fold dilutions of DNA template from HHV-6 strains were tested using quantU32 F-R and species-specific origin primers (Table S1) using cycling conditions of 95C 30s and 40 cycles of 95C 5s and 60C 30s.For the U95 deletion in the CO strains, CO4 142970F-143228R primers (Table S1) were used in the same cycling conditions with SsoAdvanced Universal SYBR Green SuperMix, while the U32-targeting qPCR was performed with the U32 primers and a Taqman probe (Table S1) with the same cycling conditions.

### PCR and Agilent Tapestation analysis

PCR across the origin tandem repeat was performed using 1 ng of template genomic DNA in 20uL total volume reactions using 10pmol of each primer and the Phusion High-Fidelity DNA polymerase according to manufacturer’s instructions. PCR primer sequences can be found in Table S1. PCR reactions were analyzed with the Genomic DNA ScreenTape assay on an Agilent 4200 TapeStation.

### Amplification-free nanopore sequencing

Nanopore libraries were created using the SQK-RAD002 kit tagmentation library preparation with 100ng of input total genomic DNA from the HHV-6B Z29 strain cultured in SupT1 cells. Amplification-free tagmented libraries were run according to Oxford Nanopore protocols v1.3.24 on a singular Mk1 (R9.4) FLO-MIN106 flow cell. Nanopore reads were mapped to the HHV-6 Z29 reference genome (NC_000898) using the LASTZ and Geneious aligners to screen for origin-containing reads (26, 27).

## Results

### Large tandem repeats covering the origin of replication in both HHV-6A and HHV-6B strains

In order to obtain single nucleotide resolution and copy number measurement for HHV-6 type strain reference materials, we sequenced a HHV-6A GS strain obtained from the HHV-6 Foundation and a HHV-6B Z29 strain obtained from the NIH AIDS repository. Libraries from the HHV-6 Z29 and GS strain were each prepared twice and sequenced to an average depth of 76X and 453X, respectively. The HHV-6B Z29 strain contained a homogeneous 983 bp long tandem repeat (Figure 1A). Copy number estimates based on relative coverage at the edge of the repeats across multiple library preparations indicated an average of 11-13 copies of the repeat present. Mapping of the edges of the Z29 origin tandem repeat gave different repeat breakpoints than previously described (28). All Z29 strains sequenced in this study had an additional 123 nucleotides at the 5’ end of the repeat and extra 4 nucleotides at the 3’ end of the repeat than the previously described repeat to make the 983 bp tandem repeat (28).

**Figure 1.**
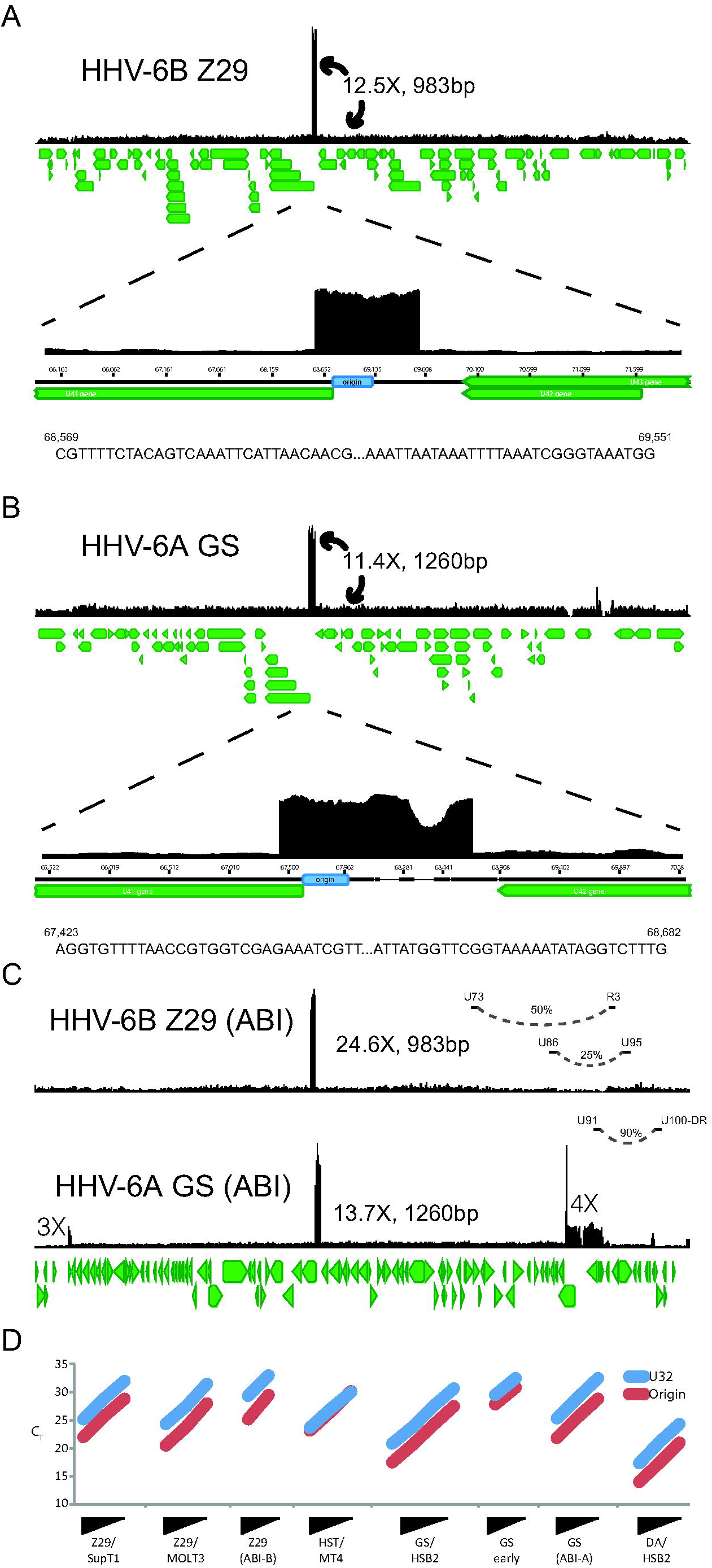
Representative coverage maps of HHV-6B Z29 and HHV-6A GS reference strains. Shotgun DNA sequencing reads from cultured virus were mapped to the NCBI HHV-6 reference genomes, NC_000898 and NC_001664, respectively. A) HHV-6B Z29 strain yielded a homogeneous 983 bp tandem repeat that was present at approximately 12.5X higher coverage of the rest of genome. Sequences at 5’ and 3’ end of the tandem repeat in Z29 strain are depicted and are different than those indicted in original paper (28). B) HHV-6A GS strain yielded a heterogeneous 1254 bp tandem repeat that was present at approximately 11.4X higher coverage than the rest of the genome. Sequences at 5’ and 3’ end of the heterogeneous tandem repeat in GS strain are depicted. C) ABI Quantitative DNA material for HHV-6A GS and HHV-6B Z29 also demonstrates a similar origin tandem repeat with additional loci with copy number differences in the GS strain. Long-distance rearrangements between U73-R3, U86-U95, and U91-U100/DR intergenic region are noted by arching dotted lines and the estimated viral subpopulation containing the indicated deletion is indicated by the % value. D) qPCR analysis of U32 and origin loci on 10-fold dilutions of DNA from HHV-6A and 6B strains confirms deep sequencing estimates of relative copy number. Equivalent Cts for the PCR results of HST/MT4 indicate an equal amount of amplification from the 2 sites with that strain. All other strains show increased amplification with the Origin PCR, indicating additional copies of the origin present in the genome. Of note, an early passage GS strain showed 3-fold less amplification of the origin by qPCR compared to the later passage GS-HSB2 strain.

HHV-6A GS strain contained a heterogeneous tandem repeat that covered 1260 bp of the HHV-6A reference genome (Figure 1B). Copy number estimates based on relative coverage at the edge of the repeats indicated an average of between 10-12 copies of the repeat present. The most common tandem repeat present included deletions of 193 bp and 2bp along with an insertion of 189 bp based on the HHV-6A reference genome, giving a mode length of 1254 bp.

To demonstrate that the copy number heterogeneity we found in the type strains of HHV-6 are also present in commercially available quantitative clinical reference materials, we also performed shotgun DNA sequencing on an HHV-6A GS strain and HHV-6B Z29 strain from Advanced Biotechnology Inc (ABI). These strains had similar tandem repeats in size at the origin of replication present in the GS strain obtained from the HHV-6 Foundation and Z29 strains obtained from NIH AIDS repository (Figure 1C). However, the Z29 origin tandem repeat in the commercial reference material was present at approximately twice the copy number observed in Z29 from the NIH AIDS repository. Intriguingly, the HHV-6A GS strain quantitative secondary standard material also contained a 4-fold increase in coverage covering the U90, U91, and the N-terminal two-thirds of the U86 gene. The U91 end of the repeat contained a complex rearrangement with the U100-DR intergenic region 266 nucleotides 5’ of the beginning of the annotated DR repeat region. The HHV-6B Z29 secondary standard contained two large rearrangements -- one between U73 and R3 repeat region constituting 50% of DNA present and another between U86 and U95 representing 25% of DNA present. Thus, copy number of HHV-6B between U86 to R3 was 4-fold lower and between U73-U86 and R3-U95 was 2-fold lower than the rest of the genome. Quantitative PCR analysis of origin and U32 loci confirmed deep sequencing data, demonstrating a 3-4 cycle earlier Ct for origin tandem repeat containing strains compared to strains lacking origin tandem repeats. Of note, an early passage GS strain had 3-fold fewer copies of the origin than the later passage GS-HSB2 strain, consistent with what has been described previously in Z29 (28).

Previous analysis of 125 HHV-6B genomes obtained from clinical specimens revealed no tandem repeats across the origin of replication. To confirm the origin tandem repeat present in the HHV-6B Z29 strain, we performed PCR amplification and fragment analysis by gel electrophoresis across the sequence tandem repeat with two separate primer sets. As a control, we performed PCR across the origin of replication in a HHV-6B PCR-positive patient specimen. The patient specimen demonstrated a single copy of the locus present, while the Z29 strain contained an amplification ladder consistent with a population of virus with different numbers of multiple tandem repeats present at the locus (Figure 2A). We also performed amplification-free nanopore sequencing on DNA extracted from the HHV-6B Z29 strain in culture. Across 6,369 nanopore reads, we recovered two reads that contained more than one copy of the origin tandem repeat (Figure 2B). One read contained three tandem repeats of the origin repeat, while the other contained two repeats. Neither read spanned the entire tandem repeat.

**Figure 2.**
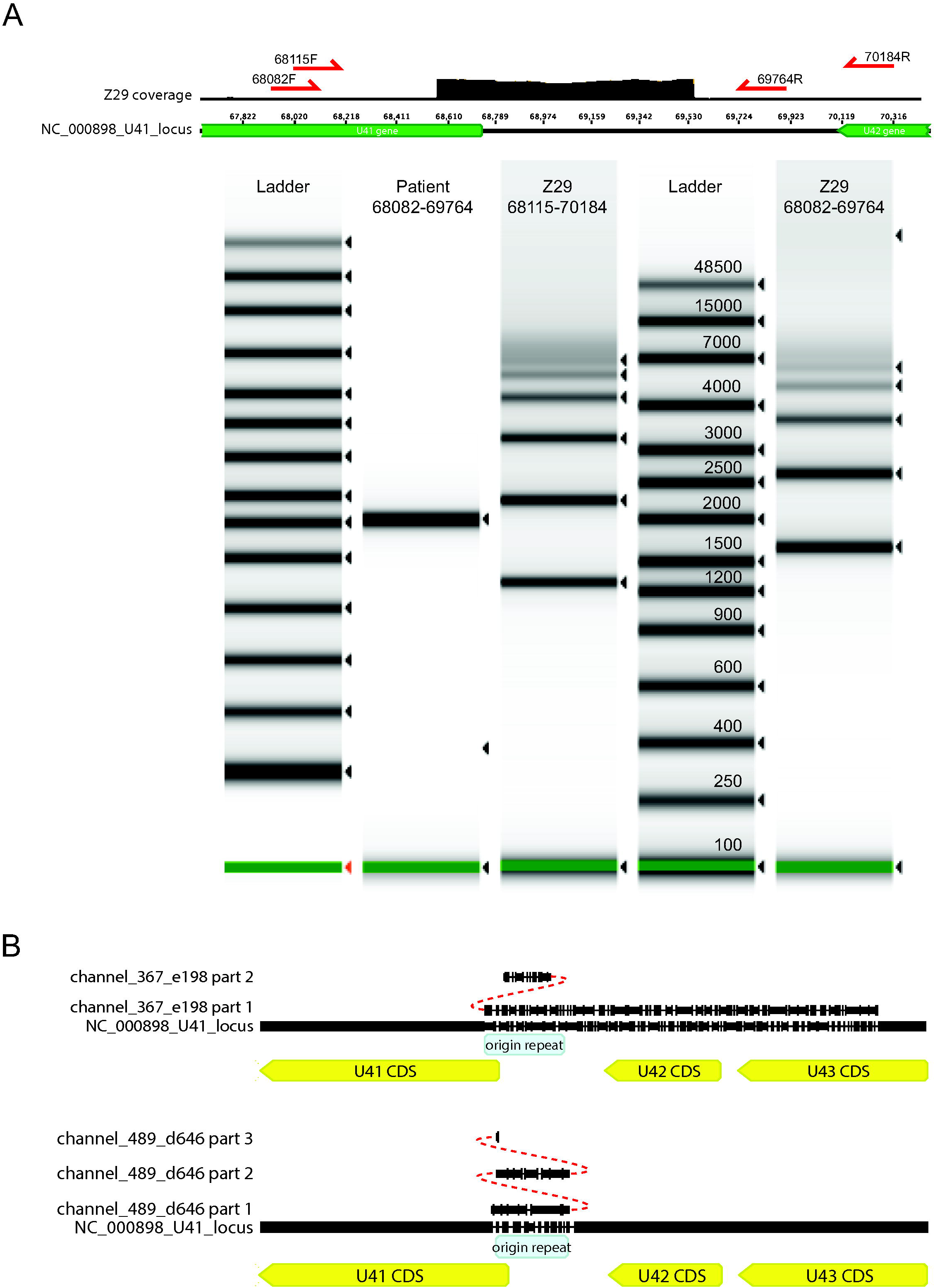
Validation of Z29 origin tandem repeat with PCR and nanopore sequencing.A) PCR-TapeStation analysis of tandem repeat in origin of Z29 strain with two PCR primer pairs. B) Amplification free-nanopore sequencing yielded two nanopore reads that align across the origin tandem repeat and carry at least 2 and 3 copies of the repeat. No reads that spanned both ends of the tandem repeat were recovered.

### Interspecies recombination between HHV-6A and HHV-6B in strain DA

Whole genome sequencing of the HHV6 DA strain revealed a hybrid genome indicative of interspecies recombination between HHV-6A and HHV-6B strains. The DA strain genome overall showed closest sequence identity to HHV-6A than HHV-6B strains but included an oriLyt repeat that measured the exact length of the Z29 repeat at 983 bp (Figure 3A). The DA strain U38 gene matched with perfect identity to the HHV-6A GS strain by BLASTN analysis. Of note, the DA strain origin-binding protein U73 gene matched closer to the HHV-6A U73 than HHV-6B U73 (99.5% versus 97.1% pairwise nucleotide identity to HHV-6A and HHV-6B reference strain genomes, respectively).

**Figure 3.**
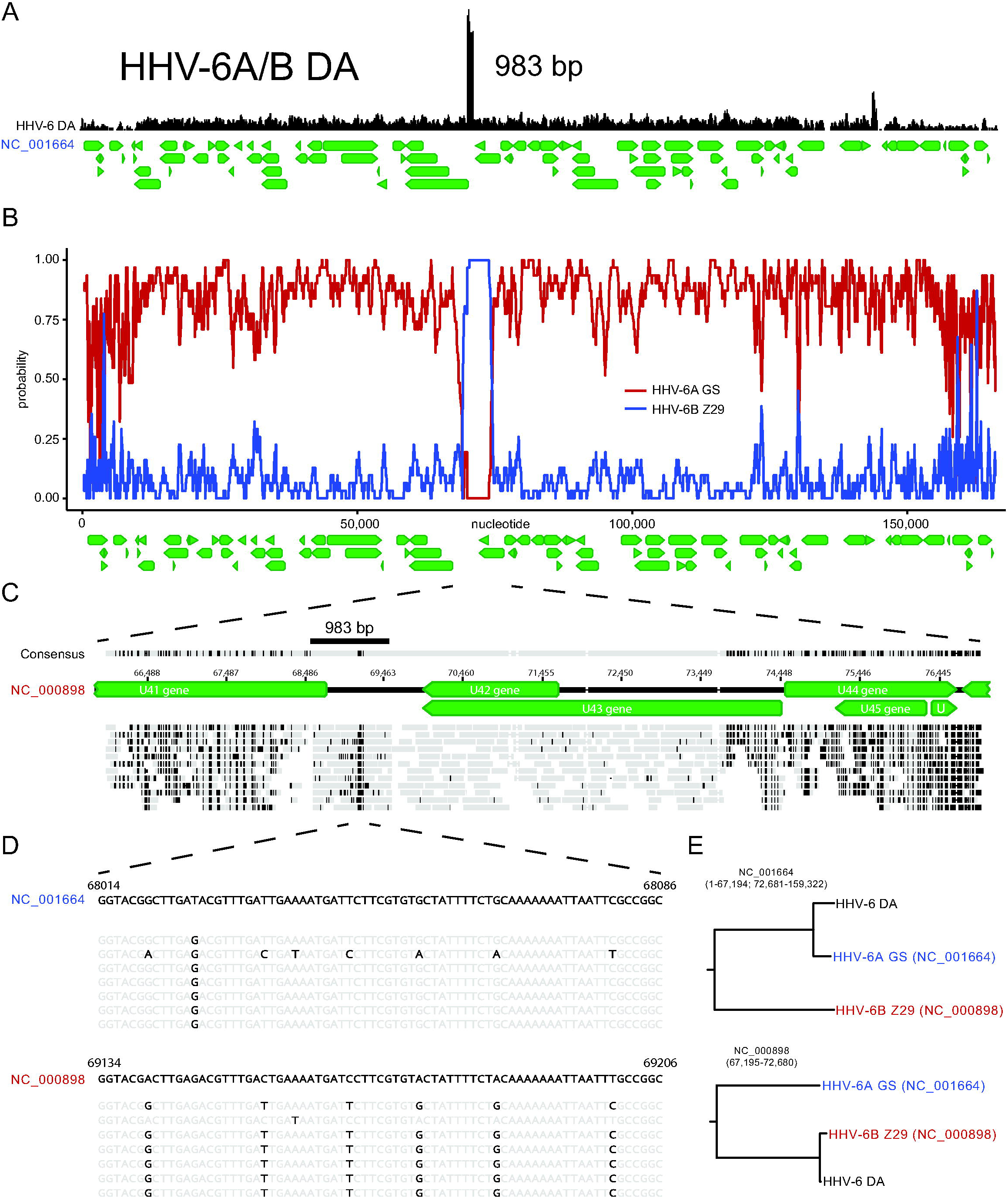
HHV-6A DA strain shows genomic evidence of interspecies recombination between HHV-6A and HHV-6B strains. A) Tandem repeat of the origin for DA strain shows Z29-like length of 983 bp. B) RDP4 scan recombination analysis demonstrates two breakpoints recombination breakpoints at nucleotide 67,194 and 72,681 of the HHV-6A DA genome C) Mapping of reads to HHV-6B Z29 reference genome at U41 locus with mismatches highlighted. D) Nucleotide sequence of 61bp region of tandem repeat that most closely matches HHV-6A reference genome (NC_001664). E) Phylogenetic tree analysis of recombination region supports HHV-6B-like nature of U41 locus.

Analysis of the DA strain sequence yielded consensus recombination breakpoints at 67,314 bp and 72,667 bp of the NC_001664 that were detected by six of seven recombination analysis programs (RDP4, GENECONV, Bootscan, MaxChi, Chimaera, and 3Seq) (Figure 3B). Reads mapped with near identity along a 5.5kb region of the HHV-6B Z29 reference genome between the 5’ end of U41 and 3’ end of U43 (68535-73824 bp of NC_000898) (Figure 3C). Only a 61bp fragment with oriLyt repeat had 6 variant sites to HHV-6B sequences and matched identically to the HHV-6A sequences in this (Figure 3D). These sequences were just 3’ from the end of the minimal origin of DNA replication annotated in the HHV-6A reference genome. The length of the HHV-6B sequence present in the DA strain is likely considerably longer than the 5.5kb shown on the reference genome, due to the presence of the oriLyt repeat. Strain DA also contained a 33bp deletion in the 3’ end of U79 gene that is not represented in either HHV-6A or HHV-6B reference sequences.

### Large deletions in U12-U24 genes and U94-U95 genes from two laboratory-adapted HHV-6A strains from collagen vascular disease

Five HHV-6A strains isolates from patients with collagen vascular disease also grew to high copy number in HSB2 cells. Isolates CO1, CO2, and CO3 were cultured in primary peripheral blood lymphocytes for 17-21 days and then HSB2 cells for 4-5 days, while isolates CO4 and CO7 were cultured for 2 days in primary cells and 42-46 days in HSB2 cells (22). The five CO isolates were highly similar with an average pairwise nucleotide identity of >99.9%, while all five isolates most closely aligned to HHV-6A isolate GS (KJ123690.1, 99.2% pairwise nucleotide identity). Both isolates CO4 and CO7 had 60% decreased coverage in a 13.5kb region from U12-U24 relative to the CO1-CO3 isolates (Figure 4A). Most notably, both isolates CO4 and CO7 also had 95% lower coverage over a 4.9kb region covering genes U94 and U95 relative to CO1-CO3 isolates. Equivalent relative copy number estimates for both the U95 locus and the origin of replication were recovered by qPCR (Figure 4B/C). Isolates CO4 and CO7 both had equivalent mixed variant allele frequencies at 25 loci based on read mapping to the UL region of the HHV-6A reference genome (NC_001664) (Table S2). No other variants were isolated between the HHV-6 CO4 and CO7 strains, suggesting these strains are identical.

**Figure 4.**
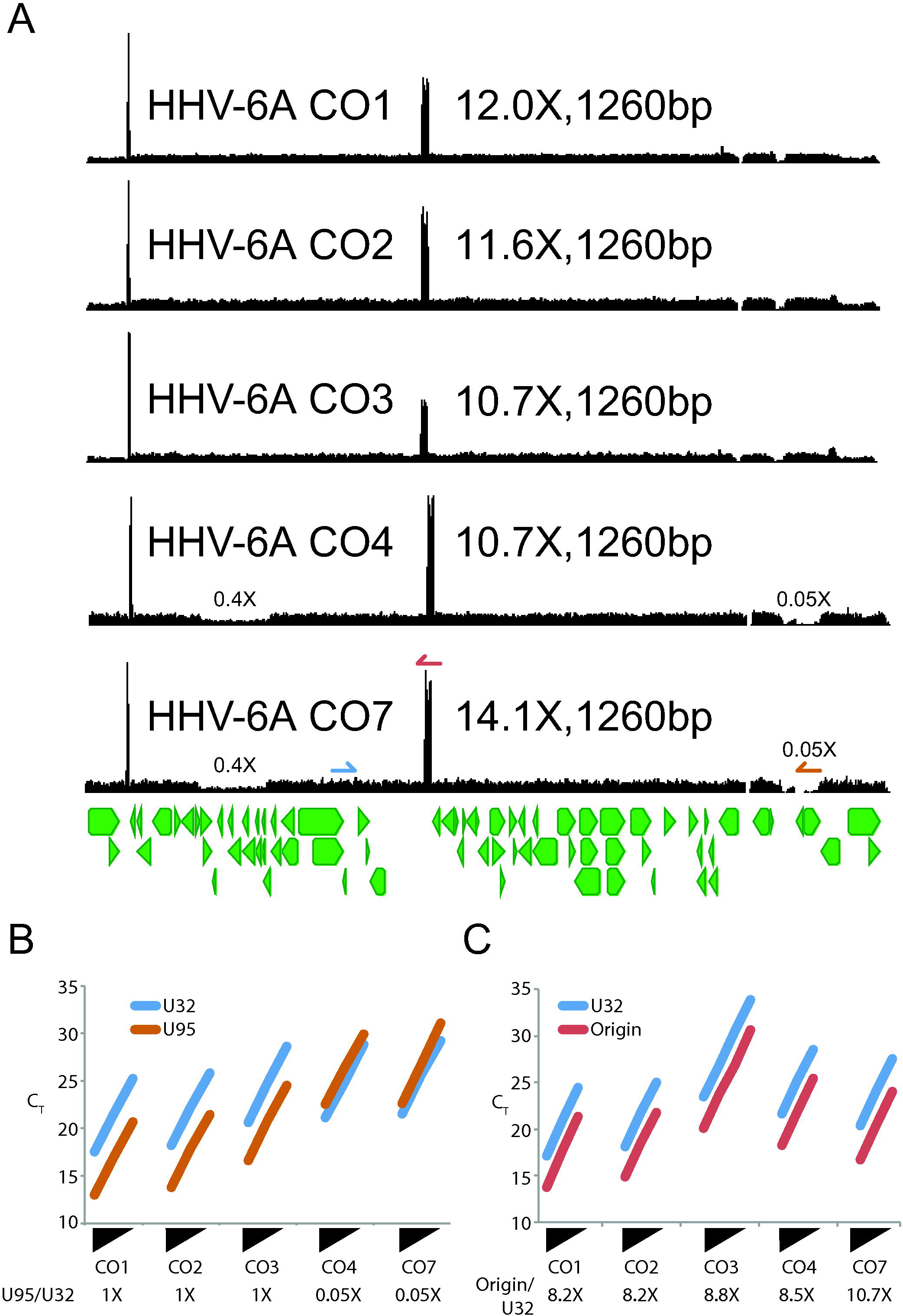
HHV-6A CO strains from patients with collagen vascular diseases show several copy number differences. A) Coverage maps from five HHV-6A strains isolated from different patients with collagen vascular diseases are displayed (22). These strains all showed a similar heterogeneous tandem repeat that gave a mode length of 1254 bp. Strains CO4 and CO7 which were passaged >40 times in immortalized HSB2 cell lines also demonstrated 60% and 95% lower coverage in U12-U24 and U94-U95 regions, respectively. Of note, these two strains also shared identical sequence and minor allele distribution, consistent with being the same strain. B/C) qPCR analysis of ten-fold dilutions of DNA from the HHV-6A CO strains at the U32, U95, and origin loci confirms relative copy number estimates from deep sequencing data.

### HHV-6 strains isolated from PHA-stimulated cord blood mononuclear cells do not contain large tandem duplications or deletions

To find the best secondary standard for HHV-6 clinical testing and to better understand the origin of the origin tandem repeat, we sequenced four HHV-6B and one HHV-6A strain that were isolated from PHA-stimulated cord blood mononuclear cells. Four HHV-6B strains (HST, ENO, KYO, NAK) were isolated from Japanese exanthem subitum patients, one HHV-6B strain (MAR) was isolated from an asymptomatic French child, and one HHV-6A strain (SIE) was isolated from an Ivory Coast patient with adult T-cell leukemia (23, 24, 29). Interestingly, all six strains contained minimal levels of copy number heterogeneity with an average coefficient of variation of coverage in the unique long region of 21.0%, compared with 119% averaged over HHV-6A GS and HHV-6B Z29 type strains (Figure 5A). The lack of origin amplification in these CBMC-passaged strains was also confirmed by qPCR (Figure 5B). Without the tandem repeat in the origin, the only source of copy number differences in the cord blood mononuclear passaged strains was direct repeat coverage at two-thirds of that in the unique long region, consistent with active replication and sequencing of mostly HHV-6 concatemers (Figure 5A) (30). Of note, the decreased coverage in the direct repeat region relative to the unique long region was present in all strains sequenced in this study.

**Figure 5.**
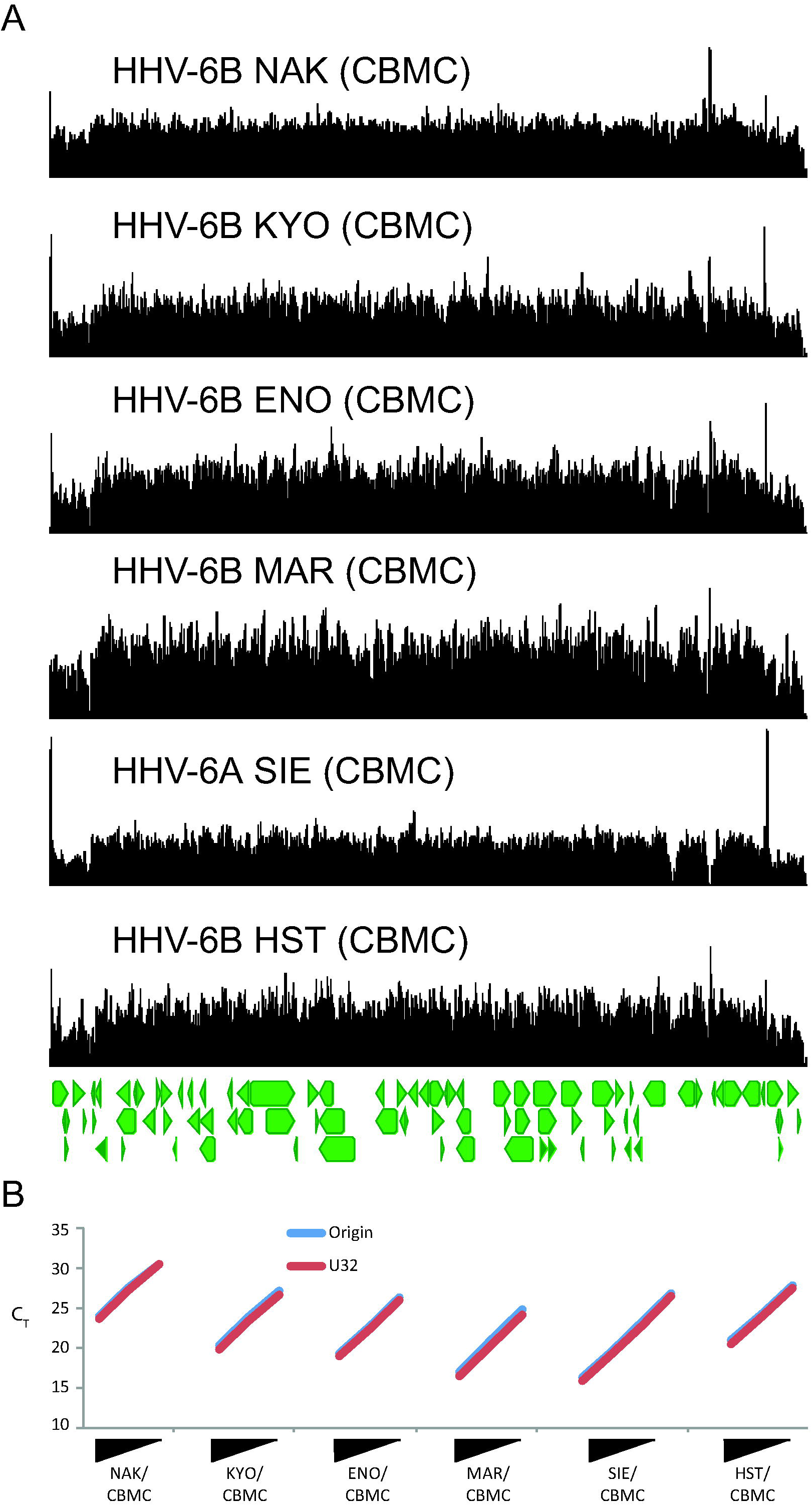
HHV-6A and -6B strains sequenced from PHA-activated cord blood mononuclear cells reveal no major copy number differences. Coverage maps from four HHV-6B strains from Japan (NAK, KYO, ENO, HST), one HHV-6B strain from France (MAR), and one HHV-6A strain from Ivory Coast (SIE) are displayed. The only copy number difference in these strains is the reduced coverage in the direct repeat region due to sequencing of likely concatemeric HHV-6, which was present in all strain sequenced in this study. B) qPCR analysis of ten-fold dilutions of DNA from the CBMC passaged HHV-6 strains at the U32 and origin loci confirms relative copy number estimates from deep sequencing data.

## Discussion

We show the presence of copy number heterogeneity at multiple genomic loci across HHV-6A and HHV-6B culture isolates that are used as reference material for clinical assay development and normalization as well as basic science virology work. The most prominent copy number difference was a large tandem repeat in the origin of replication that was present in both HHV-6A and HHV-6B strains. The presence of large copy number differences in HHV-6A and HHV-6B strains was associated with passaged in immortalized cell lines, as strains passaged in primary cell lines and cord blood mononuclear cells did not carry genomic duplications and deletions and early passage virus contained fewer tandem repeats than late passage virus.

Recent analysis of 130 HHV-6B genomes and 10 iciHHV-6A sequences spanning the U41 origin region revealed no large tandem repeats in the origin of replication among clinical isolates (18). Previous work had illustrated a large heterogeneous tandem repeat present at the oriLyt in HHV-6B Z29 strains that was associated with higher passage number (28). Our data illustrate that this is a general feature of HHV-6 and that a heterogeneous larger tandem repeat is also present in multiple laboratory adapted HHV-6A strains at the oriLyt. We show that increased culture passage in immortalized cell lines is associated with reduced copy number of two large genomic loci in HHV-6A CO strains. Copy number variability at multiple genomic loci was also reflected in HHV-6A reference material that is used to normalize quantitative values for clinical assay development. Interestingly, no HHV-6 isolate from cord blood mononuclear cells showed tandem repeats, despite growing to high titer. These isolates may provide the best material for HHV-6 standards development.

We also show the first genomic evidence of interspecies recombination between HHV-6A and HHV-6B strains along a >5.5 kb segment containing the oriLyt and U42 gene as well as portions of the U41 and U43 genes. Interspecies recombination is a relatively common feature of the alphaherpesviruses HSV-1 and HSV-2 but has not been described for any other human herpesviruses (31). Our results for DA strain recombination are most consistent with a model in which co-cultivation of an HHV-6A GS strain with a laboratory-adapted HHV-6B Z29 strain with oriLyt repeat that resulted in recombination between the two strains. Interestingly the recombinant strain also showed HHV-6A-like sequence in its HHV-6B oriLyt tandem repeat. Although this sequence fell outside the minimal origin of replication, it is suggestive that the oriLyt tandem repeat sequence may have evolved to more effectively interact with HHV-6A replication proteins contained in the rest of genome. No specific HHV-6B-like sequences were found in the DA strain U73 origin binding protein to indicate reciprocal U73 evolution to match the HHV-6B-like origin sequences. Recent analysis of 140 HHV-6 genomes from clinical and iciHHV-6 isolates revealed no evidence of interspecies recombination but widespread intraspecies recombination (18). Whether interspecies recombination has occurred in clinical strains of HHV-6 remains to be determined.However the high sequence similarity between HHV-6A and HHV-6B and the frequency of alphaherpesvirus recombination suggests that as more sequencing is performed, HHV-6A and HHV-6B recombinants may be found in nature.

Previous work from our group had demonstrated the loss of almost one-third of the BK and JC polyomavirus genome in up to 90% of viral species present in multiple viral stocks including a WHO international standard, likely due to viral passage in SV40 T-antigen immortalized cell lines (19, 20). While clinical PCR tests for HHV-6 are unlikely to target the oriLyt region, normalization of quantitative clinical HHV-6 testing to any of the loci here found at increased or decreased copy number could affect quantitation, not least because multiple viral populations were present in many of the reference materials tested. We also found several examples where contamination with another HHV-6 strain most parsimoniously explained our sequencing results. We continue to recommend the use of next-generation sequencing to obtain genome-wide single nucleotide resolution copy number measurements in order to validate viral reference materials used in clinical virology and basic sciences labs across the world.

Accessions: These sequences are available in NCBI Genbank (MF994813-MF994829) and associated with BioProject 338014.

